# A multi-network comparative analysis of transcriptome and translatome in cardiac remodeling

**DOI:** 10.1101/2020.07.01.181743

**Authors:** Etienne Boileau, Shirin Doroudgar, Eva Riechert, Lonny Jürgensen, Thanh Cao Ho, Hugo A Katus, Mirko Völkers, Christoph Dieterich

## Abstract

Our understanding of the transition from physiological to pathological cardiac hypertrophy remains elusive and largely based on reductionist hypotheses. Here, we profiled the translatomes of 15 mouse hearts to provide a molecular blueprint of altered gene networks in early cardiac remodeling. Using co-expression analysis, we reveal how sub-networks are orchestrated into functional modules associated with pathological phenotypes. We show how transcriptome networks are only partially reproducible at the translatome level. We find unappreciated hub genes and genes in the transcriptional network that were rewired in the translational network, and associated with semantically different subsets of enriched functional terms, providing novel insights into the complexity of the organization of *in vivo* cardiac regulatory networks.

## 1 INTRODUCTION

Exercise- and disease-induced cardiac growth are associated with different molecular profiles and differ in the signalling pathways that drive remodeling; yet both are characterized by an increase in size of cardiomyocytes, sarcomerogenesis, and overall increase in heart-weight-to-body-weight (HW/BW) ratio (Shimizu and Minamino, 2016). While adaptive exercise-induced hypertrophy allows the heart to maintain an adequate cardiac output with improved contractility, pathological hypertrophy is a maladaptive response, concurring with irreversible changes (*e*.*g*.. cardiomyocyte loss, fibrosis, reduced cardiac function), and typically progressing to heart failure (Shimizu and Minamino, 2016; Bernardo et al., 2018).

Little is known about the molecular mechanisms controlling physiological hypertrophy, particularly from a multiomics systems biology perspective. Yet our understanding of the *in vivo* transition from adaptive hypertrophy to cardiac dysfunction has important clinical implications (Boström et al., 2010). There has been substantial interest in characterizing genes and regulatory pathways that underpin the disease phenotypes, and the implications of transcriptional reprogramming and translational regulation at various stages of the hypertrophic response. Only recently have post-transcriptional regulatory networks been uncovered that are of central importance for morphological remodeling in fibrosis (Chothani et al., 2019), or for modulating the early reponse to cardiac stress (Doroudgar et al., 2019).

In this study, we adopt a systems biology approach to integrate multi-omics data through the use of co-expression networks to highlight higher-order relationships among gene programs that are expressed in the heart *in vivo* under growth stimuli. In such networks, genes are connected if there is a significant co-expression relationship between them (Langfelder and Horvath, 2008). Modules or sub-networks represent clusters of genes with related function or involved in common processes or pathways. Our analysis of 15 mouse left ventricular tissues from experimental models of exercise- and disease-induced cardiac hypertrophy showed, for the first time, the organization of the transcriptome and translatome into networks of biologically meaningful clusters of co-expressed genes. By correlating module expression and disease phenotypes, we were able to show the synchronized expression dynamics of genes encoding extracellular matrix, and cytoskeletal proteins, and a diminished contribution of electron transport complex genes, genes associated with oxidative phosphorylation and mitochondrial function. In contrast to differential expression analysis, co-expression and rewiring analysis led us to the identification of yet uncharacterized candidate genes, key to organizing the behaviour of transcriptome and translatome networks. Our results also reveal how transcriptome networks are only partially reproducible at the translatome level. Our findings provide a molecular blueprint of altered gene networks during the early phases of *in vivo* cardiac remodeling.

## 2 RESULTS

### 2.1 Exercise-*vs*. disease-induced cardiac remodeling

To characterize stress-induced cardiac remodeling and the response to acute and chronic pressure overload, we used the swim model (Wang et al., 2010) and the transverse aortic constriction (TAC) model. The transverse aortic constriction is a commonly used model to mimic pressure overload-induced cardiac hypertrophy and heart failure (Rockman et al., 1991). The swim model is referred to as the physiological or the healthy model, and represents exercise-induced hypertrophy. The TAC model is referred to as the pathological model, and represents disease-induced hypertrophy.

To study co-expression network models of translational regulation during cardiac remodeling, we used *in vivo* Ribo-seq and RNA-seq libraries from 15 mouse hearts representing experimental models of physiological and pathological cardiac hypertrophy. We used the RiboTag approach to capture the cardiac translatome. Cardiomyocyte-specific analysis of ribosome protected fragments was achieved after affinity purification using the RiboTag mouse (Doroudgar et al., 2019) (Methods, and Fig. 1 A). To catalog translation events in the mouse heart, we performed an unsupervised search for actively translated open-reading frames (ORFs) using Rp-Bp (Malone et al., 2017) (Fig. S1, and Methods ‘Detecting active translation’). The final list of translated genes was used as background for co-expression and differential expression analyses (Methods ‘Constructing gene co-expression networks’, and ‘Differential expression and translational efficiency analysis’). Co-expression networks were constructed separately for RNA-seq (transcriptome) and Ribo-seq (translatome) count data and used to identify hub genes associated with cardiac remodeling. All translated ORFs, RNA-seq and Ribo-seq read counts can be found as supporting information (Table EV1, Table EV2).

**FIGURE 1.**
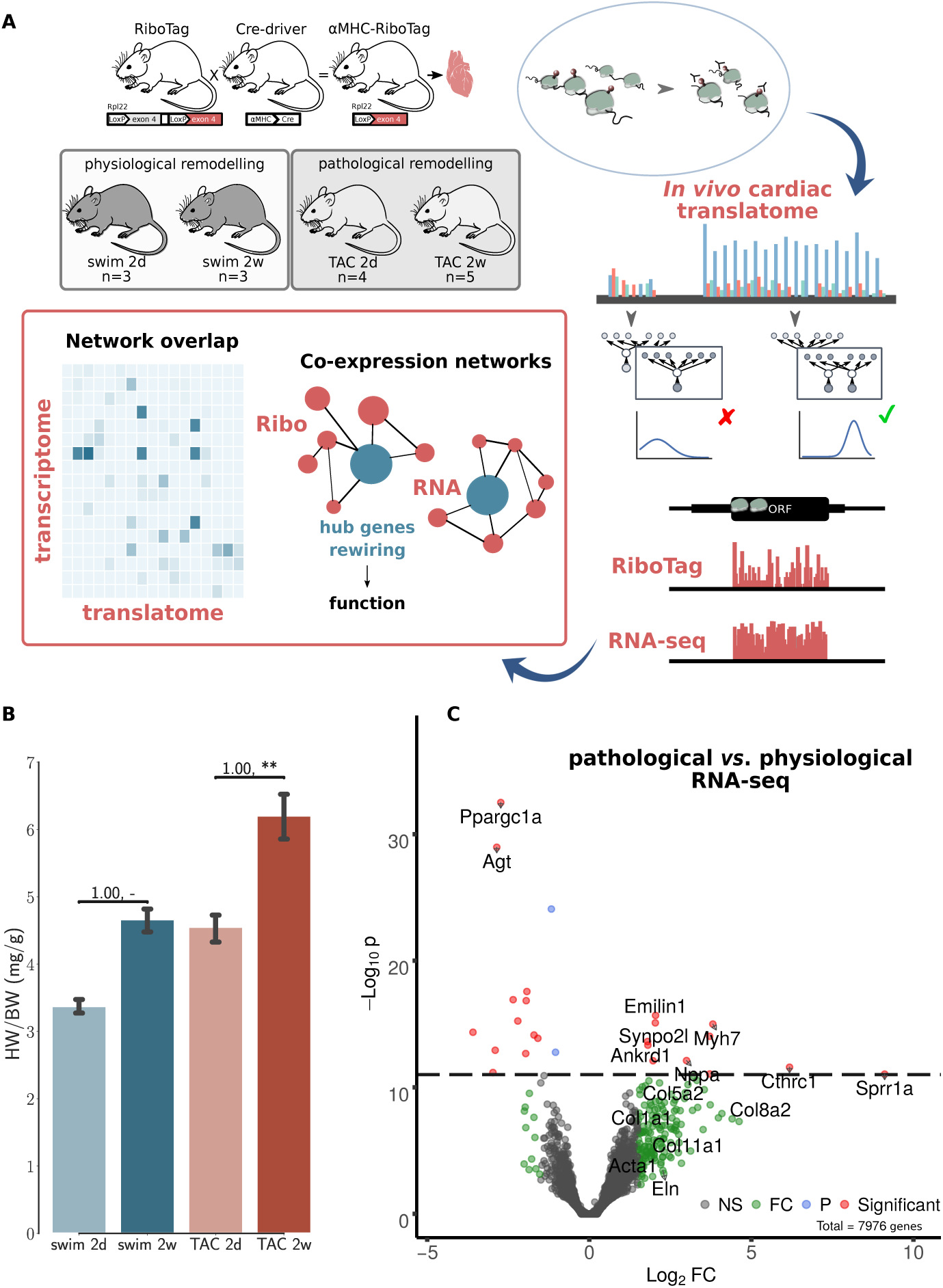
Co-expression networks of transcriptional and translational regulation in cardiac remodeling. (A) The RiboTag system is used to find the cardiac translatome of 15 animals from two models at two time points. A final set of translated genes is inferred using graphical Bayesian models. Co-expression networks are then constructed separetely for *in vivo* Ribo-seq (translatome from RiboTag data) and RNA-seq (transcriptome) data. The networks were used to identify hub genes associated with cardiac remodeling, and how changes in connectivity (network rewiring) may be associated with differential functionality in transcriptome *vs*. translatome. (B) Heart-weight-to-body-weight ratio (HW/BW) after 2d and 2w of exercise in the physiological model, 2d and 2w after transverse aortic constriction (TAC) surgery. N≥3 for each time point. Significance was measured using Mann-Witney U-test, at a threshold of 0.05 (** = <0.01, p-values may be affected by the small sample size, for this reason we also report effect size using non-directional rank-biserial correlation). Kruskall-Wallis test H=10.9, p-value=0.01. (C) Volcano plot displaying the distribution of all genes with relative abundance (log_2_ RNA-seq pathological *vs*. physiological) plotted against significance level, showing significantly increased and decreased genes during *in vivo* pathological stress. Top genes (red) are highlighted. A number of genes (green) significant at a threshold of 0.05 are also highlighted. NS=Non significant, FC=Fold change only significant, P=P-value only significant, Significant=FC+P.

We monitored the acute response at an intermediate time point (2 days after TAC), and a chronic time point (2 weeks after TAC), when cellular and molecular remodeling has occurred, but cardiac function is preserved (Doroudgar et al., 2019). Matching time points were monitored in the physiological model (swim at 2 days and 2 weeks). Increased HW/BW ratios were observed after 2 weeks in the swim and TAC models, with a larger increase in the pathological model (Fig. 1 B). At the RNA-seq level only, we observed an upregulation of the genes for atrial natriuretic peptide (Nppa), the fetal isoform of myosin heavy chain (Myh7) (Taegtmeyer et al., 2010), clinically relevant genes such as Ankrd1 or Synpo2l (Ling et al., 2017; van Eldik et al., 2017), as well as a number of genes implicated in tissue remodeling (Fig. 1 C). Taken together, these results are consistent with graded, pathological cardiac hypertrophy in the TAC model. In the physiological model, we did not observe fetal gene re-expression, typically associated with metabolic remodeling in a variety of pathophysiologic conditions (Taegtmeyer et al., 2010). There was a significant increase in Ppargc1a, a master regulator of mitochondrial biogenesis associated with physiological hypertrophy (Boström et al., 2010), and Agt, key component of the renin-angiotensin system (RAS), suggesting that the swim model did not induce a pathological hypertrophy phenotype, but instead improved the cellular energetics of the heart.

### 2.2 Co-expression networks of cardiac remodeling

We calculated topological overlap and clustered genes, identifying 17 distinct co-expression modules for each of the RNA-seq and Ribo-seq networks (Table EV3). The modules were labeled in order from RNA1 to RNA17, and from Ribo1 to Ribo17, using unsupervised hierarchical clustering based on co-expression correlation with disease association (Fig. 2 A, Fig. 3 A). Our analysis revealed how gene expression programs in the heart are organized differently in transcriptome and translatome space into modules, or sub-networks, of highly connected genes. Gene Ontology (GO) enrichment analysis suggest that significant genes in a number of modules localize to common cellular components, such as the extracellular matrix (ECM) and associated proteins (RNA1, RNA7, Ribo1, Ribo2, and Ribo4), the cytoskeleton, related membrane ruffling (RNA4 and RNA6) and the cell cortex (Ribo8, Ribo10), the Golgi apparatus (RNA2), the nucleosome (Ribo12), or the various sub-compartments of the mitochondrion (RNA13, RNA14, RNA15, Ribo13, and Ribo16) (Table EV4). These modules have one or more related molecular function or are associated with shared biological processes.

**FIGURE 2.**
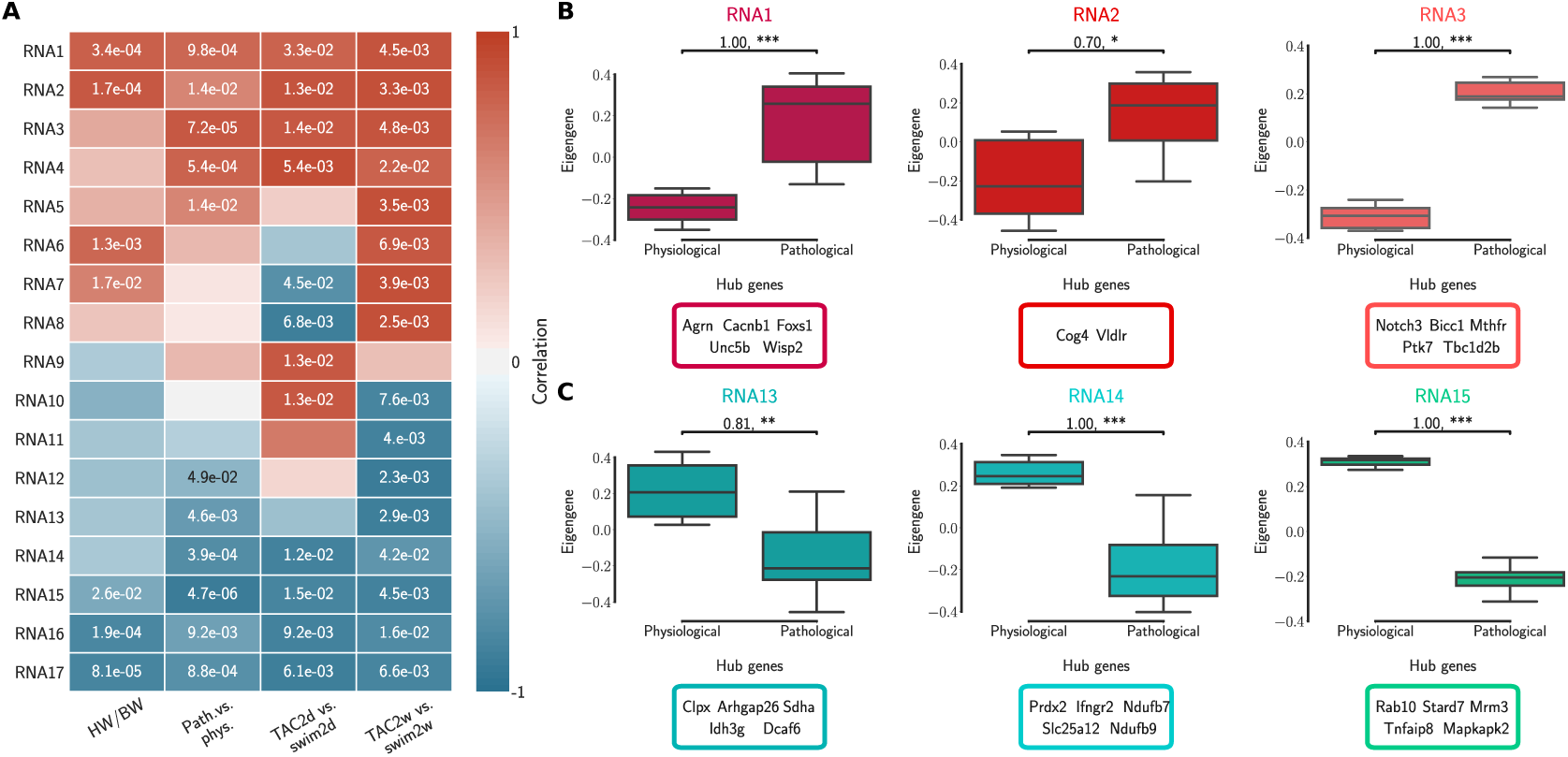
Transcriptome (RNA) network modules were clustered to assess relatedness based on correlation of co-expressed eigengenes. (A) Biweight midcorrelation and Student correlation p-values between eigengene cluster expression and disease association as well as heart-weight-to-body-weight ratio. (B) The first three cluster expression profiles and top ranked hub genes that are positively correlated with pathological cardiac remodeling. (C) The first three cluster expression profiles and top ranked hub genes that are negatively correlated with pathological cardiac remodeling (*i*.*e*. positively correlated with the physiological model). Significance was measured using a one-sided Mann-Witney U-test, at a threshold of 0.05 (** = <0.01, ***= <0.001, p-values may be affected by the small sample size, for this reason we also report effect size using non-directional rank-biserial correlation). Up to five hub genes are shown.

**FIGURE 3.**
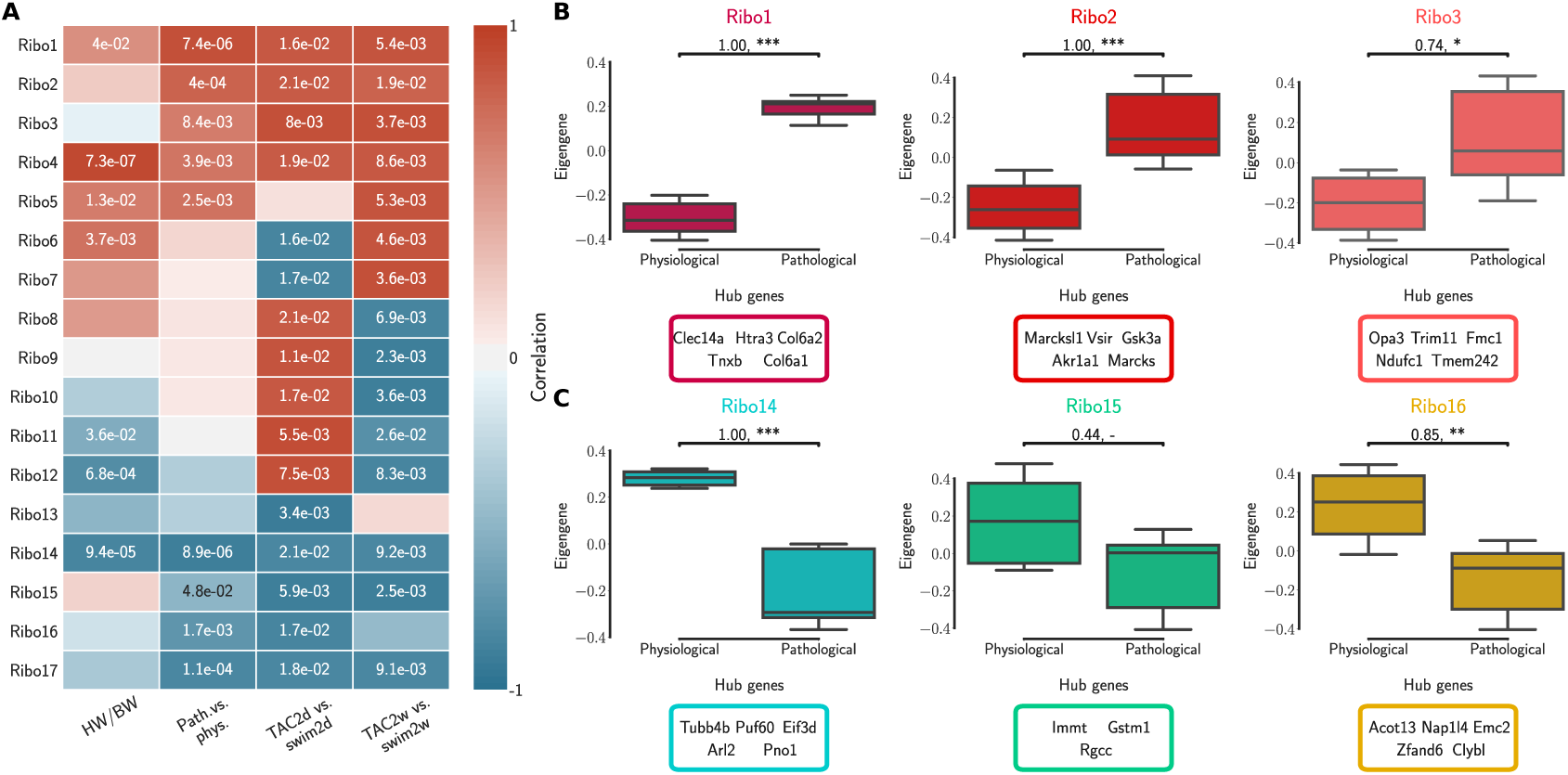
Translatome (Ribo) network modules were clustered to assess relatedness based on correlation of co-expressed eigengenes. (A) Biweight midcorrelation and Student correlation p-values between eigengene cluster expression and disease association as well as heart-weight-to-body-weight ratio. (B) The first three cluster expression profiles and top ranked hub genes that are positively correlated with pathological cardiac remodeling. (C) The first three cluster expression profiles and top ranked hub genes that are negatively correlated with pathological cardiac remodeling (*i*.*e*. positively correlated with the physiological model). Significance was measured using a one-sided Mann-Witney U-test, at a threshold of 0.05 (** = <0.01, ***= <0.001, p-values may be affected by the small sample size, for this reason we also report effect size using non-directional rank-biserial correlation). Up to five hub genes are shown.

#### 2.2.1 Correlation with the cardiac pathophysiology of remodeling

We calculated the module correlations to disease association and HW/BW ratios, and clustered them to identify sub-networks associated with patho-physiological features and pre-clinical symptoms of cardiac hypertrophy. Five RNA modules and five Ribo modules had a positive correlation to the pathological model and to increased HW/BW ratio, and almost all were significant (RNA1 to RNA5, Ribo1 to Ribo5, Fig. 2 A, and Fig. 3 A). We also observed a negative correlation to the pathological model (*i*.*e*. a positive correlation to the physiological model) in five RNA modules and, to a moderate extent, in four Ribo modules (RNA13 to RNA17, Ribo14 to Ribo17). RNA1 to RNA5 (Ribo1 to Ribo5) were activated after pressure overload in the pathological model, whereas RNA13 to RNA17 (Ribo14 to Ribo17) were repressed, when looking at the module eigengenes (Fig. 2 B, C, Fig. 3 B, C, and Fig. S2 A, B, Fig. S3 A, B). An eigengene is a representative of the standardized module expression values across all samples. Eigengenes have been largely regarded as robust biomarkers (Oldham et al., 2008; Johnson et al., 2018; Zhang et al., 2018; Di et al., 2019). The strong association between RNA1 to RNA5 (Ribo1 to Ribo5), on the one hand, and that of RNA13 to RNA17 (Ribo14 to Ribo17), on the other hand, suggests a synchronized expression dynamics characterized by an increased role of genes encoding ECM and cytoskeletal proteins, and a diminished or altered contribution from mitochondrial translation, metabolic pathways of carbohydrate, fat, and protein metabolism, as well as oxidative phosphorylation. RNA6 to RNA12 (Ribo6 to Ribo13) showed a dynamic association pattern to either the 2d or the 2w time points, uncovering the transcriptional and translational heterogeneity in response to pressure overload (Fig 2 A, Fig 3 A, and Fig. S2 C, Fig. S3 C). The significance of these associations was highly consistent, and when we looked at the correlation between module membership and gene significance, we found that the strongest and most significant associations were for the top five modules, particularly for RNA (RNA1 to RNA5) (Table EV5). In addition, for modules RNA6 to RNA12, gene significance at 2w was more often and more strongly correlated with module membership, suggesting that genes associated with the later time points are the key drivers. On the contrary, for most modules Ribo6 to Ribo13, the association was observed with the earlier time points, suggesting a higher relative contribution of translational control at 2d, consistent with a rapid translational response to stress (Doroudgar et al., 2019).

A hallmark of pathological hypertrophy is the reactivation of fetal gene expression (Taegtmeyer et al., 2010). Key markers of the fetal gene program were found in sub-networks correlated with the pathological models (Table EV3): Nppa (RNA1, Ribo1), Nppb (RNA3, Ribo7), or Myh7 (RNA3, Ribo1). While Myh6 was found in RNA15 and Ribo16, consistent with the observed known ‘gene switches’, we observed the presence of several other genes clustered in RNA1, Ribo4 or Ribo2 (Myh7b, Myh10, Myh11, and Myh14), and whose clinical significance has not yet been described in the context of cardiac hypertrophy. These switches were also observed for Glut1 and Glut4: Slc2a8 (RNA1, Ribo1), Slc2a1 (RNA10, Ribo10), Slc2a4 (RNA14, Ribo3), or Slc2a12 (RNA15, Ribo17); and for Myc: Mycn (RNA1, Ribo4), and Myc (RNA17, Ribo10).

In each module, we also identified highly connected intramodular hub genes, which may function as core regulators of the hypertrophic response (Fig. 2 B, C, Fig. 3 B, C, and Fig. S2, Fig. S3). Intramodular connectivity measures how co-expressed is a given gene with respect to all other genes of a particular module. Among the top candidates of modules one to five (RNA and Ribo), we found a large number of structural genes encoding for ECM proteoglycans, collagen, fibrillar, and cell adhesion proteins, as well as protein filaments. In RNA13 to RNA17 (Ribo14 to Ribo17), many of the most highly connected hub genes had a strong support of mitochondrial localization.

#### 2.2.2 Co-expression networks uncover hub genes not found by differential expression

Unsupervised hierarchical clustering based on hub gene expression showed that the top interacting genes serve as a molecular signature to differentiate physiological and pathological models of cardiac hypertrophy (Fig. 4 A, and Fig. S4 A). These observations prompted us to verify whether hub genes were also differentially connected (DC) between RNA-seq and Ribo-seq networks, and how these observations compare with results of differential translational-efficiency (DTE) analyses. Up to 80% of DC genes were unchanged in DTE, but less than 5% of these were hub genes (Fig. S5). Although DC could highlight variations that are not reflected by differences in gene expression or translational efficiency, these results suggest that the most important genes in each cluster remain relatively unchanged in DC when comparing RNA-seq and Ribo-seq.

**FIGURE 4.**
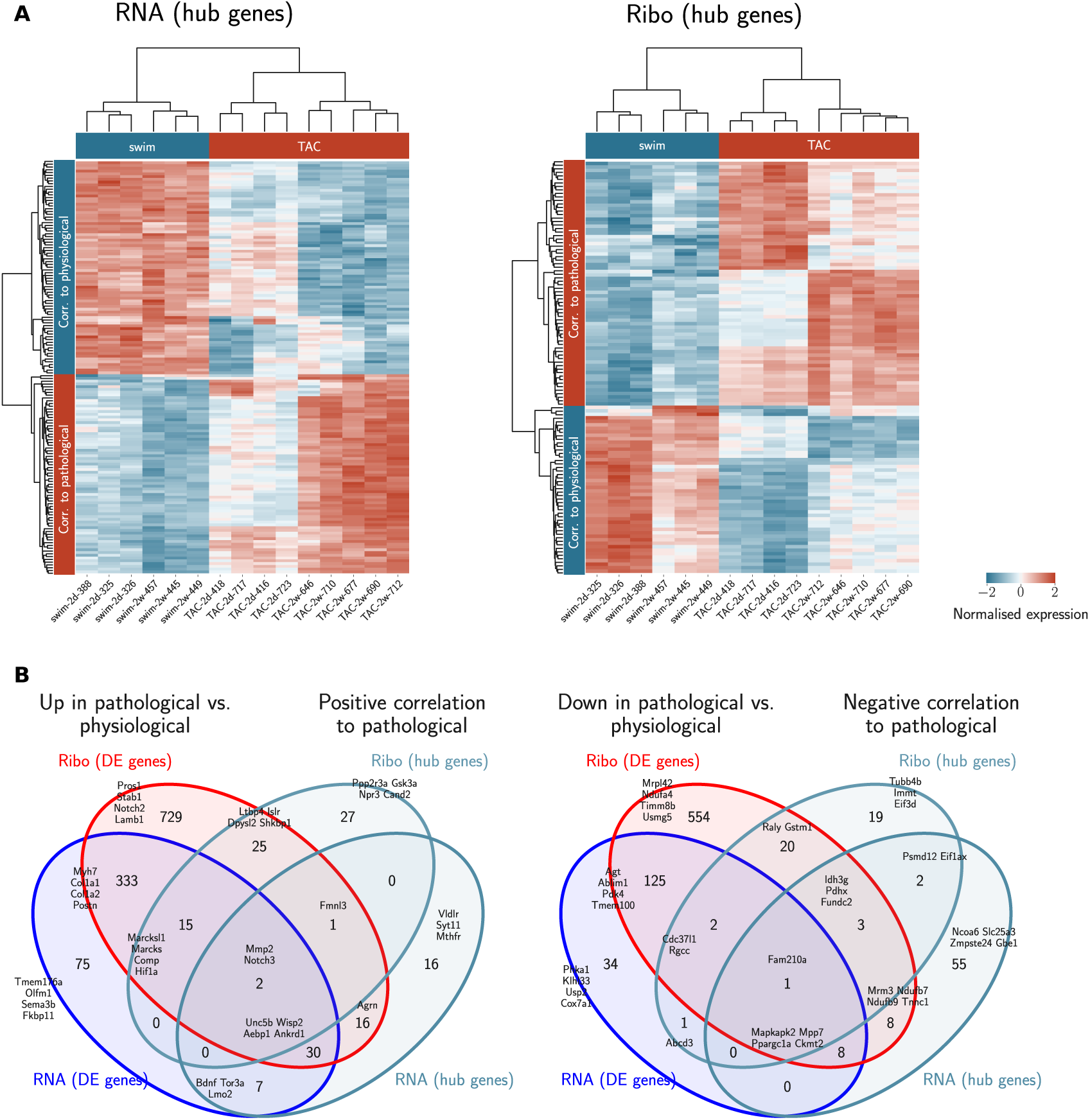
Hub genes provide a different molecular signature for differentiating physiological and pathological cardiac remodeling. (A) Expression profile of all hub genes associated with co-expression clusters having a positive or a negative correlation to pathological phenotypes after unsupervised hierarchical clustering, showing a separation between samples of the physiological and pathological models, and between positively and negatively correlated eigengene-associated cluster. Hubs genes are from clusters RNA1 to RNA5, RNA13 to RNA17, and from Ribo1 to Ribo5, Ribo14 to Ribo17. (B) *Left* Venn diagram showing the overlap between differentially up-regulated genes at the transcriptional and translational level in the pathological model, and hub genes in the RNA-seq and Ribo-seq co-expression clusters with positive correlation to pathological phenotypes. *Right* Venn diagram showing the overlap between differentially down-regulated genes at the transcriptional and translational level in the pathological model, and hub genes in the RNA-seq and Ribo-seq co-expression sub-networks with negative correlation to pathological phenotypes. Transcriptionally-regulated genes are DTE in RNA, or RNA+Ribo; translationally-regulated genes are DTE in Ribo, RNA+Ribo, or TE (FDR < 0.05, fold change > log_2_(1.2)). Top four differentially regulated genes are indicated. For all subsets intersecting hub genes, up to four selected hub genes among the top ranked genes (<15) are also indicated. DE=Differentially expressed.

Many of the top hub genes of modules with positive correlation (RNA1 to RNA5, and Ribo1 to Ribo5) were also up-regulated in the pathological model (Fig. 4 C). Similarly, a number of top hub genes of modules with negative correlation (RNA13 to RNA17, and Ribo14 to Ribo17) were down-regulated in the pathological model. Similar observations were made considering the different time points and the varying correlations, using hub genes from modules RNA6 to RNA12, and Ribo6 to Ribo13 (Fig. S4 B). While the number of differentially regulated genes is much larger, we found hub genes from co-expression correlation only that were not identified in DTE, and whose significance in the context of cardiac injury and remodeling is discussed below.

#### 2.2.3 Co-expression networks describe the organization of the heart transcriptome and translatome

To extract an additional layer of regulatory information, we investigated the degree of preservation between RNA network structure and Ribo co-expression network, and the amount of overlap between sub-networks. We identified modules that were highly correlated/anti-correlated with the pathological model that were partially shared across transcriptome and translatome (RNA1, RNA3, and RNA4 overlap with Ribo1, Ribo2 and Ribo4; RNA14, RNA15, and RNA17 overlap with Ribo14, Ribo16, and Ribo17). These could represent common and continuous mechanisms of response to stress (Fig. 5 A). Among all these, five (six) RNA clusters were found to be highly (moderately) preserved at the translatome level (Fig. 5 B). Preservation is based on density and connectivity measures, and uses the correlation structure of the networks to identify differences between RNA-seq and Ribo-seq. Six more RNA modules, one of which was activated (RNA5) after pressure overload, in the pathological model, two of which were repressed (RNA13 and RNA16), as well as RNA7, RNA8, and RNA11, which showed differential activation/repression at 2d and 2w, had no or little gene overlap with translatome modules, and did not have a preserved network structure, suggesting that the transcriptome does not capture all key changes occurring in the heart during early hypertrophy.

**FIGURE 5.**
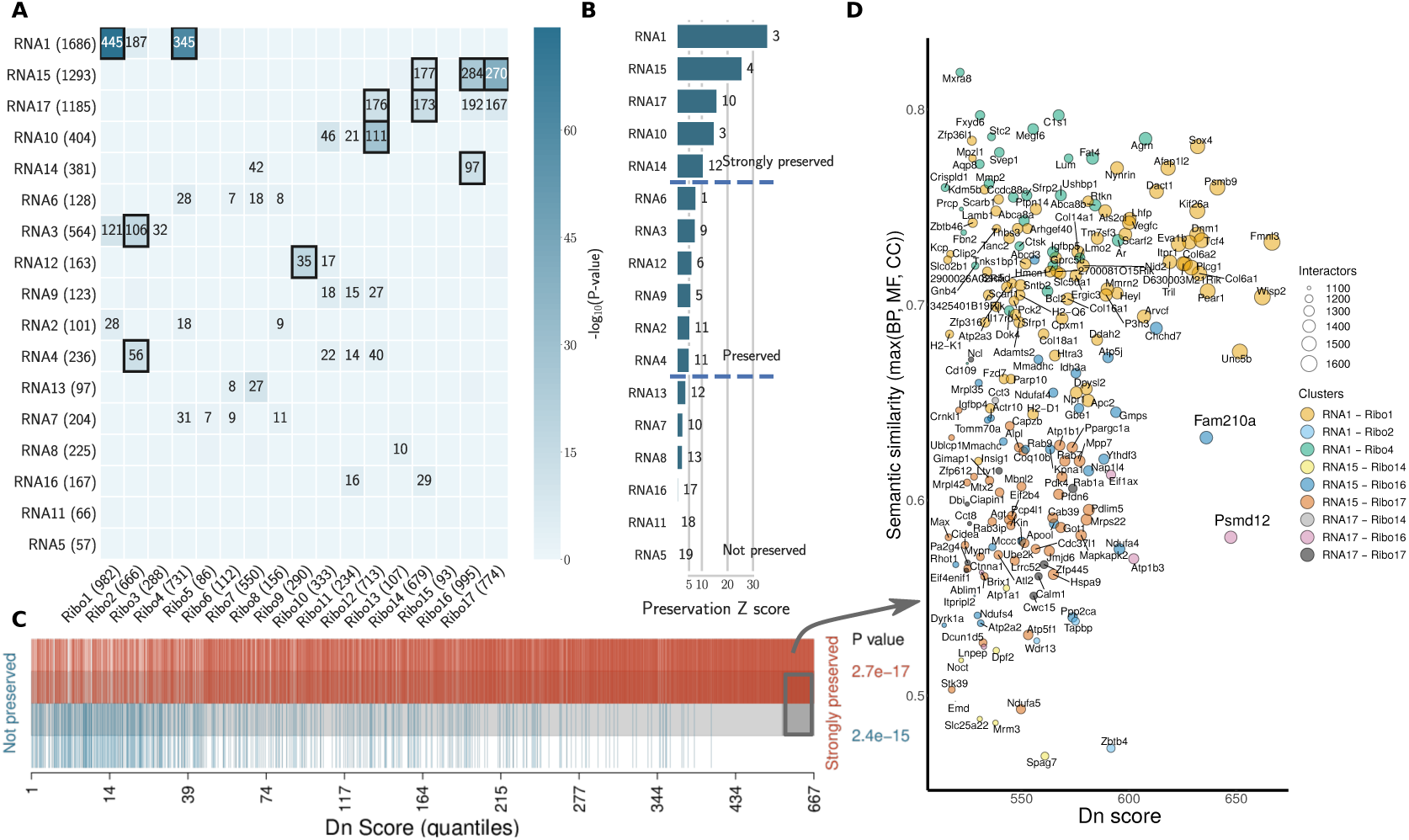
The organization of the heart transcriptome and translatome under cardiac remodeling. (A) Cross-tabulation of RNA-seq (rows) and Ribo-seq (columns) clusters. Each row and column is labeled by the corresponding cluster and its size (number of genes). Counts of genes in the intersection is shown for clusters sharing a significant overlap (p-value < 0.01, two-tailed Fisher’s exact test). Overlap with p-value < 1e-10 are highlighted. (B) Preservation Z scores (mean of *Z* -scores computed for density and connectivity measures) for the RNA-seq co-expression clusters in the Ribo-seq network. The vertical lines indicate the threshold for moderate preservation (5) and strong preservation (10). The *medi an Rank* statistic (mean of the observed density and connectivity statistics) is indicated to the right of each bar. (C) Barcode plot showing the enrichment of genes based on their dynamic neighborhoods score (Dn score), compared to their association with clusters that are either strongly or not preserved. Genes are ranked according to their Dn score, and colored based on their association. The grey box towards the right indicates genes for which Dn score is greater than two standard deviations above the average. To test whether genes belonging to preserved or not preserved clusters are highly ranked in terms of Dn score, the camera test from the limma R package was performed and p-values are shown. (D) Scatter plot of the most rewired genes (grey box in C). A semantic similarity analysis was performed for each ontology (BP, MF, and CC) and every gene using its immediate interactors in the RNA-seq and Ribo-seq networks, respectively, and the maximum semantic similarity is reported on the y-axis. The set of all translated genes was used as background. Dots are colored according to their cluster association, and the size represents the total number of interactors, or connected genes, in a hypothetical joint network combining the respective RNA-seq and Ribo-seq clusters. Semantic similarity was calculated using the GOSemSim R package. Only genes in clusters with significant overlap are reported. BP=Biological Process, MF=Molecular Function, CC=Cellular Component.

To uncover how the heart transcriptome and translatome networks are rewired in response to stress, we highlighted genes which had the most dynamic neighborhoods (Methods, ‘Dynamic neighborhoods score’). The Dn score, or dynamic neighborhoods score, captures changes in connectivity of a gene, even when its intramodular connectivity remains similar in RNA-seq and Ribo-seq networks, and is thus better suited than DC to highlight hub genes associated with potential regulatory mechanisms. Interestingly, the most rewired genes all belonged to the top preserved modules (Fig. 5 C). Among these, we highlighted two candidates, Fam210a and Psmd12, identifed earlier (Fig. 4), which were strongly rewired, and whose immediate interactors were enriched in different GO terms between RNA-seq and Ribo-seq networks (Fig. 5 D, and Fig. S6). Fam210a is a conserved transmembrane protein localized in the mitochondria, containing a mitochondrial targeting signal peptide (MitoCarta2.0 mouse), a DUF1279 (Domain of Unknown Function) domain with a transmembrane peptide, and a coiled coil at the C-terminus (InterPro). It is mostly expressed in the heart [1.72534] and skeletal muscle [1.57137] (Standardized values, BioGPS Mouse Cell Type and Tissue Gene Expression Profiles), and is thought to play a role in modulating muscle and bone biology (Tanaka et al., 2018), but its function in the heart and its molecular mechanisms are unknown. Psmd12, encoding the non-ATPase regulatory subunit 12 of the 26S proteasome, is better characterized. The 26S proteasome is a multiprotein complex involved in the ATP-dependent degradation of ubiquitinated proteins, and thus plays a key role in protein homeostasis. Psmd12 is associated with several pathways, including Regulation of Apoptosis, Stabilization of p53, and p53-Dependent/Independent G1 DNA Damage Response (WikiPathways, Reactome). The tumor suppressor Trp53 (p53) regulates cell growth and fate, and its role in the heart is well-known. Psmd12 is also associated with inflammation [2.13579] and hypertrophy [2.10896] (Standardized values, CTD Comparative Toxicogenomics Database). Overall, module-eigengene association to disease phenotypes has led to the identification of highly rewired hub genes that may function as drivers of cardiac remodeling. These hub genes are potentially involved in related, but different molecular pathways or functions, suggesting some form of translational control that may not be immediately apparent from DTE analyses.

## 3 DISCUSSION

In this study, a model of left ventricular pressure overload was used to mimic hypertrophy induced by systemic hypertension and aortic stenosis, and compared with a physiological model of exercise-induced cardiac growth. Transcriptional and translational co-expression networks uncovered *in vivo* changes in the heart occuring within two weeks of a transverse aortic constriction (TAC) surgery, revealing the complexity of the organization as well as unappreciated genes that may act as key drivers of the hypertrophic response.

Physiologic and pathophysiologic stimuli act upon the cell membrane and work their way through various cascades to mediate gene expression, translational control and protein levels (Haque and Wang, 2017). As expected, pressure overload was associated with profound changes in the composition of the extracellular matrix (ECM), which were reflected by a sub-clustering and a synchronized expression dynamics of ECM-, and cytoskeletal-related genes, in both transcriptome and translatome networks (Figs. 2, S2, 3, and S3, Table EV3). A marked upregulation of genes encoding ECM proteins has previously been observed during the transition from stable cardiac hypertrophy to heart failure (Boluyt et al., 1994). Our results indicate that concerted dynamic changes occur early *in vivo* after stimuli, and are likely to be implicated in transducing molecular signals driving the maladaptive response. Concurrently to these observations, modules anti-correlated to the pathological model showed a diminished expression or altered contribution of mitochondrial, electron transport complex and oxidative phosphorylation genes. These modules (1-4, and 14, 15, 17) were also among the most preserved (Fig. 5), suggesting that some critical sub-network structures remain relatively stable, although genes associated with these may or may not show significant changes in translational/transcriptional efficiency (Fig. 4 and S4).

Co-expression network and differential translational-efficiency (DTE) analyses are based on different assumptions (Langfelder and Horvath, 2008, 2012). In co-expression networks, the top genes are the most connected genes, based on the correlation structure. We identified a number of hub genes that may function as molecular drivers of cardiac remodeling: Gsk3a (Ribo2), characterized for its role in regulating glycogen metabolism, and shown to be a critical player in cardiac hypertrophy (Sugden et al., 2008; Zhou et al., 2016); Cand2 (Ribo2), a mTOR dependent mRNA recently identified as a growth regulator, upregulated in the acute phase of cardiac hypertrophy (Sandmann et al., 2018); Vldlr (RNA2), characterized for its role in obesity associated lipotoxicity (Fungwe et al., 2019), and negatively associated with survival in the ischemic heart (Perman et al., 2011); Eif3d (Ribo14), thought to be involved in cap-dependent translation during cellular stress, independently of Eif4e (Lee et al., 2016); Immt (Ribo15), a mitochondrial membrane protein whose transcript is regulated by Rbm20 (Zahr and Jaalouk, 2018); Ncoa6 (RNA16), whose dysfunction is linked to dilated cardiomyopathy in mouse models (Roh et al., 2014); Slc25a3 (RNA14), the absence of which was shown to drive a novel model of metabolic, mitochondrial-driven cardiomyopathy (Kwong et al., 2014); or Zmpste24 (RNA17), which may be implicated in prelamin A toxicity-driven inflammation (Brayson et al., 2019). Among the hub genes associated with varying correlations at 2d and 2w, we found Rptor (Ribo9), encoding a component of the mTORC1 complex, pivotal for cell growth (Shende et al., 2011); Lonp1 (Ribo9), crucial for the maintenance of mitochondrial homeostasis, recently identified as a potential target in protecting the myocardium from oxidative damage (Venkatesh et al., 2019); Ubr4 (Ribo9), also recently found to play a role *in vivo* in myofiber hypertrophy in mice (Hunt et al., 2019); C5ar1 (RNA10), a mediator of a novel evolutionarily conserved response promoting cardiomyocyte proliferation after cardiac injury (Natarajan et al., 2018); S100a4 (RNA10), whose expression protects cardiomyocytes against ischemic stress (Doroudgar et al., 2016), Phb2 (Ribo13), a conserved protein located in the mitochondrial inner membrane, whose function until recently has remained largely unknown, which has recently been linked to cardiac fatty acid metabolism and heart failure (Wu et al., 2020); Ace (RNA7), recognised as a top candidate gene for cardiovascular research (Niu et al., 2002); or Sparc (RNA8), long-associated with cardiovascular risk and dysfunction, and recently characterized for its novel inotropic function in the heart (Kos and Wilding, 2010; Vaughan et al., 2018; Deckx et al., 2019).

We identified a number of hub genes in the transcriptional network that were rewired in the translational network, and associated with semantically different subsets of enriched terms (Fig. 5, S6). Notably, we highlighted the presence of two hub genes that were rewired under hypertrophic stimuli, Fam210a (RNA15, Ribo16), and Psmd12 (RNA17, Ribo16). Fam210a, a gene of previously unknown function, has been described as a musculoskeletal modulator (Tanaka et al., 2018). In humans, a prior study reported that Fam210a (C18orf19) was the strongest candidate partner protein of Atad3a (ATPase Family AAA Domain Containing 3A), which was also found in the same modules (RNA15 and/or Ribo16), along with 60 (out of 153) interacting proteins identified by Orbitrap MS analysis and quantified by SILAC labelling (He et al., 2012). Atad3a is essential for mitochondrial metabolism and translation, and has been implicated in several processes in mitochondria. Taken together, these results suggest that Fam210a could modulate translation of mitochondrial-encoded electron-transport chain proteins, and play a yet undescribed role in cardiac muscle adaptation and growth.

The biological importance of Psmd12 as a scaffolding subunit in proteasome function has been described earlier in the context of neuronal development, but remains un-documented in the heart. The ubiquitin proteasome system (UPS) is critical in preventing accumulation of damaged and misfolded proteins, and has been implicated in a number of cardiac proteinopathies and heart failure (Pagan et al., 2013; Cacciapuoti, 2014; Maejima, 2020). Our results support the existence of transcriptional/translational regulatory processes affecting the or affected by proteasome function in the pathogenesis of cardiac hypertrophy.

In summary, these results highlight the organization of distinct molecular processes into sub-networks of co-expressed genes, and describe how transcriptome and translatome signatures are orchestrated into functional modules and hub genes associated with the early stages of cardiac remodeling. We found that the higher-order organization of gene co-expression networks in the transcriptome is only partially reproducible at the translatome level, pointing to the role of post-transcriptional control as an additional layer regulating the hypertrophic response. Our results constitute a valuable resource to study *in vivo* cardiac regulatory networks, and a first step towards the identification and characterization of novel proteins involved in cardiac remodeling, hypertrophy and heart failure.

## 4 MATERIALS AND METHODS

### 4.1 Experimental models

All raw data used in this study were acquired from experimental models of pathological cardiac hypertrophy (transverse aortic constriction or TAC) and swimming-induced physiological hypertrophy. TAC (27 gauge needle) surgery was performed as previously described (Völkers et al., 2013), and animals were sacrificed after 2 days (n=4) and 2 weeks (n=5). For exercise training in the physiological hypertrophy model, mice swam regularly in a water tank for either 2 days (n=3) or 2 weeks (n=3) (Evangelista et al., 2003). The experiments were performed in 9-week-old male mice using the RiboTag system (Kmietczyk et al., 2019; Doroudgar et al., 2019). All animal experimental procedures were reviewed and approved by the Institutional Animal Care and Use Committees at the Animal Experiment Review Board of the government of the state of Baden-Württemberg, Germany.

### 4.2 Preparation of sequencing libraries

Mice were sacrificed, and their hearts were excised, washed in PBS containing 100 *µ*g/ml cycloheximide (CHX), and snap frozen in liquid nitrogen. Left ventricular tissue was homogenized using a tissue homogenizer in 5 volumes of ice-cold polysome buffer (20 mM Tris pH 7.4, 10 mM MgCl, 200 mM KCl, 2 mM DTT, 1% Triton X-100, 1U DNase/*µ*l) containing 100 *µ*g/ml CHX. Ribo-seq and RNA-seq libraries were prepared for each biological replicate from the identical lysate. Ribosome protected fragments (RPFs) were generated after immunoprecipitation of cardiac myocyte-specific polysomes with anti-HA magnetic beads after treating the lysate with RNase I (Ambion). Libraries were generated according to the mammalian Ribo-seq kit (Illumina). Barcodes were used to perform multiplex sequencing and create sequencing pools containing at least eight different samples and always an equal amount of both RNA and RPF libraries. Sample pools were sequenced on the HiSeq 2000 platform using 50-bp sequencing chemistry.

### 4.3 The RiboTag system

In the RiboTag mouse, the exon 4 of the Rpl22 gene is flanked by Loxp recombination sites, followed by an HA-tagged exon 4. When the RiboTag mouse is crossed to a Cre driver mouse, the Cre recombinase enzyme is activated resulting in the removal of the LoxP-flanked wild type Rpl22 exon 4 and replacement with the HA-tagged Rpl22 exon 4, which is incorporated into the ribosome particle. In mouse hearts, a cell-specific promotor (Myosin heavy chain, *α* isoform, or *α* MHC, encoded by the Myh6 gene) drives the expression of Cre which induces cardiomyocyte-specific HA-tagged ribosomes. RiboTag mice were purchased from Jackson Laboratory (JAX ID 011029) and bred to the *α* MHC-Cre mice line to obtain homozygous mice expressing Rpl22-HA in cardiomyocytes.

### 4.4 Detecting active translation

Translation prediction using Ribo-seq data was performed with Rp-Bp v2.0 (Malone et al., 2017), based on Ensembl release 96. We used evidence from uniquely mapped reads and periodic fragment lengths only. For each sample, the fragment lengths and ribosome P-site offsets were determined from a metagene analysis using the automatic Bayesian selection of read lengths and ribosome P-site offsets (BPPS). The final list of translation events includes, in addition to annotated ORFs, un-annotated ORFs with evidence of translation upstream or dowstream of annotated coding sequences. For the analyses, translation in non-coding regions as well as variants of canonical coding sequences, such as truncated, extended or internal ORFs, were discarded. We required ORFs to have a minimum length of 3 aa and more than 10 in-frame P-sites. The final list of translation events was further filtered to only include ORFs that were predicted in at least three samples and whose host gene was also annotated in APPRIS (Rodriguez et al., 2018), resulting in 9,129 unique genes.

### 4.5 Sequencing data alignment

Adapters removal and quality filtering was done with flexbar v3.0.3 (Dodt et al., 2012) using standard filtering parameters (no prior trimming for Ribo-seq). Reads with more than 1 uncalled base were not included in the output: *flexbar –max-uncalled 1 –pre-trim-left 0 –qtrim-format sanger –qtrim TAIL –qtrim-threshold 10*. Reads aligning to a custom bowtie2 v2.3.0 (Langmead and Salzberg, 2012) ribosomal index were discarded. Remaining reads were then aligned in genomic coordinates to the mouse genome (GRCm38.p6) with a splice-aware aligner, STAR v2.5.3a (Dobin et al., 2013), inserting annotations on the fly: *STAR –outSAMtype BAM SortedByCoordinate –outFilterType BySJout – outFilterMismatchNmax 1 –outFilterMismatchNoverLmax 0*.*04 –outFilterIntronMotifs RemoveNoncanonicalUnannotated –alignIntronMin 20 –alignIntronMax 100000 –sjdbOverhang 33 –seedSearchStartLmaxOverLread 0*.*5*

*–winAnchorMultimapNmax 100*. For the RNA-seq data, reads were trimmed from the 3’ end after adapter removal, such that the read length before alignment did match the maximum periodic fragment length of the corresponding Ribo-seq sample, as determined with the BPPS method. Finally, abundance estimates and read count to coding sequences were obtained using HTSeq-count (Anders et al., 2014): *htseq-count –type CDS –mode intersection-nonempty*, taking into account the strand-specific protocols.

### 4.6 Constructing gene co-expression networks

Read counts to coding sequences were used, as described above, only including genes that were considered to be translated, based on the criteria described in the ‘Detecting active translation’ section. We removed low variance genes and genes with the lowest sequencing-depth normalised average expression (first centile). From these, 7,976 genes with the highest connectivity were clustered on the basis of topological overlap (TO) to identify patterns of co-expression, using the WGCNA R package (Langfelder and Horvath, 2008, 2012). The network construction was done separately for Ribo-seq and RNA-seq data, on this common set of genes. We first applied a regularised log transformation, and corrected for batch effects, where applicable (Johnson et al., 2006).

Briefly, weighted adjacencies were defined based on signed co-expression similarity using biweight midcorrelation and a soft thresholding power of *β* =18. The scale-free topology fit index did not reach a soft-threshold of 0.85, so we chose a power value of 18 based on the number of samples, resulting in an average connectivity of 78 and 80 for RNA-seq and Ribo-seq networks, respectively. Sample clustering did not reveal abnormal grouping, counts were variance stabilised, and filtering was done based on variance and mean expression, without taking into account phenotypic information.

For each network, a reference TO matrix was first calculated. To produce robust and reproducible clusters, we then performed bootstrap-resampling (n=100) and computed the TO matrix for each of the resampled networks. In each case, resampling was done within the physiological (swim) or the pathological (TAC) group. The final consensus TO matrix was defined as the median of all scaled TO matrices, and used as input for hierarchical clustering. The consensus TO matrix can be viewed as a ‘smoothed’ version of the adjacency matrix. Network modules (subset of genes that form sub-networks) whose eigengenes were highly correlated were merged, and characterized by their eigengene expression and significance. To validate module membership, we applied post-hoc resampling (n=100) by subsetting TO of random modules matched by size with respect to the consensus TO for every module. A one-proportion Z-test was used to assess whether the mean TO of the random modules was higher than that of the module assigned by the hierarchical clustering and merging algorithm.

The module membership is defined as the correlation (biweight midcorrelation) between eigengene and gene expression values, and measures the importance of a gene within a cluster. Gene significance is defined as the correlation (biweight midcorrelation) between genes and biological traits or disease association. A gene that is significant for a given phenotypic trait may represent pathways associated with the trait. For binary/discrete variable correlation, biweight midcorrelation is replaced by the standard Pearson correlation. Hub genes were defined as genes with the highest intramodular connectivity. We ranked genes in each module and selected as hub genes those having a number of interactions greater than two standard deviations above the average connectivity found in a given module, *i*.*e*. with a Z-score > 2.

Preservation statistics were derived using the RNA-seq network as a reference, using the correlation structure of the networks. We used the *medi an Rank* statistic, defined as the mean of the observed density and connectivity statistics, and the *Z*_*summary*_, defined as the mean of *Z* -scores computed for density and connectivity measures (Langfelder et al., 2011). The *medi an Rank* was used to compare relative preservation among clusters. The *Z*_*summary*_ was used to assess the significance of observed statistics by distinguishing preserved from non-preserved clusters via permutation testing (n=100).

Differentially connectivity (DC) was defined as the 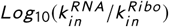. It is not a robust measure, and it is based uniquely on intramodular connectivity. A gene was DC if this ratio was greater than two standard deviations above the average across all genes.

### 4.7 Dynamic neighborhoods score

The Dn score, or dynamic neighborhoods score, of a given gene is calculated based on the variance of the state-space adjacency matrix over the network states (RNA-seq and Ribo-seq), relative to the mean centroid, as 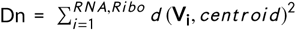, where *d* is the Euclidean distance, and **V**_**i**_ is a vector of genes in the RNA-seq and Ribo-seq networks (Goenawan et al., 2016). A dynamic neighborhood is the induced sub-graph consisting of all genes adjacent to a given gene, *i*.*e*. all immediate interactors. To calculate the Dn score, we used the TO matrix, thresholded at 0.1 (all interactions below this threshold were considered inexistant).

### 4.8 Differential expression and translational efficiency analysis

To allow for proper comparison, all RNA-seq quantifications were derived from trimmed single-end reads mapped to coding sequences, as explained in the ‘Sequencing data alignment’ section. Expression quantification and differential expression analysis was performed using DESeq2 (Love et al., 2014). The calculation of change in translational efficiency was done using an interaction term (assay + condition + assay:condition) with a likelihood-ratio test. Regulation status of a gene at the transcriptional and/or translational level can be integrated using the fold changes from standard Wald test (RNA-seq, Ribo-seq) and the likelihood-ratio test (translational efficiency). All analyses were performed on the same set of translated genes (n=7,976) used as input for network clustering. We used genome-wide significance threshold of FDR < 0.05, and a fold change (FC) of log_2_(1.2).

### 4.9 Gene ontology (GO) enrichment

GO enrichment (The Gene Ontology Consortium, 2018) to assign functional annotation to clusters of genes were performed with topGO v2.34.0 (Alexa and Rahnenfuhrer, 2018). To define the relevant gene sets corresponding to each clusters, we considered hub genes and genes with strong module membership and significance for a given trait (heart-weight-to-body-weight ratio, pathological *vs*. physiological, TAC2d *vs*. swim2d, or TAC2w *vs*. swim2w). The latter were identified by taking the upper quartile of genes with the highest module membership and gene significance for a trait having the highest correlation between absolute values of module membership and gene significance. The universe of genes consisted of all translated genes (n=9,129).

## Data availability

The data generated for this study have been deposited in NCBI’s Sequence Read Archive through the BioProject accession numbers PRJNA484227 and PRJEB29208. All raw counts and translation events used in this study are available as supporting information.

Rp-Bp is publicly available at https://github.com/dieterich-lab/rp-bp under the MIT License. The code for generating co-expression networks is available as supplementary information.

## Author contribution

EB designed the project, performed data analysis and interpretation, maintained the Rp-Bp software and wrote the manuscript. ER, LJ, TCH performed animal experiments. HAK contributed to the design of the project. SD and MV contributed to the design of the project, data acquisition and interpretation. CD conceived and designed the project, contributed to data analysis and interpretation, and helped to draft the manuscript. All authors read and approved the final manuscript.

## Acknowledgements

EB and CD acknowledge support by the Klaus Tschira Stiftung gGmbH [00.219.2013]. CD, SD and MV acknowledge the DZHK (German Centre for Cardiovascular Research) Partner Site Heidelberg/Mannhein for funding. MV is supported by the Deutsche Forschungsgemeinschaft, Plus 3 Programme of the Boehringer Ingelheim Foundation (BISI) and Baden Württemberg Stiftung. CD and EB thank the Dieterich Lab group members for providing feedback on the manuscript.

## Conflict of interest

The authors declare that they have no conflict of interest.

## Supporting information

